# Unmasking of novel mutations within *XK* gene that could be used as Diagnostic Markers to Predict McLeod syndrome: Using in silico analysis

**DOI:** 10.1101/540153

**Authors:** Mujahed I. Mustafa, Mohamed A. Hassan

**Affiliations:** Department of Biochemistry, University of Bahri, Sudan; Department of Biotechnology, Africa city of Technology, Sudan

**Author notes:** Crossponding author: Mujahed I. Mustafa.

**Keywords:** McLeod neuroacanthocytosis syndrome, X-linked recessive, central nervous systems, red blood cells, in silico analysis, XK gene, neurologists, hematologists, clinical geneticists

## Abstract

**Background:** McLeod neuroacanthocytosis syndrome is a rare X-linked recessive multisystem disorder affecting the peripheral and central nervous systems, red blood cells, and internal organs.

**Methods:** We carried out in silico analysis of structural effect of each SNP using different bioinformatics tools to predict substitution influence on protein structural and functional level.

**Result:** 2 novel mutations out of 104 nsSNPs that are found to be deleterious effect on the XK structure and function.

**Conclusion:** The present study provided a novel insight into the understanding of McLeod syndrome, SNPs occurring in coding and non-coding regions, may lead to RNA alterations and should be systematically verified. Functional studies can gain from a preliminary multi-step approach, such as the one proposed here; we prioritize SNPs for further genetic mapping studies. This will be a valuable resource for neurologists, hematologists, and clinical geneticists on this rare and debilitating disease.

## Introduction

McLeod syndrome (MSL) is one subtype of rare X-linked recessive neuroacanthocytosis syndromes, approximately 150 cases have been reported worldwide.(1) Characterized by misshapen red blood cells and progressive degeneration of the basal ganglia disease including: movement disorders, cognitive alterations, and psychiatric symptoms and cardiac manifestations. Neuromuscular symptoms leading to weakness and atrophy of different degrees.(2–6) Hematologically, MLS is defined as a specific blood group phenotype that results from absent expression of the Kx erythrocyte antigen and weakened expression of Kell blood group antigens.(3, 7) Kell antigens are encoded by the KEL gene on the long arm of chromosome 7. Kx antigen is encoded by the *XK* gene on the short arm of the X chromosome.(8)

McLeod syndrome caused by mutations of *XK* gene, an X-chromosomal gene of unknown function. But all we know is different *XK* mutations may have different effects upon the *XK* gene product and thus may account for the variable phenotype. (9–15) The *XK* gene provides instructions for producing the XK protein, which carries the blood antigen Kx. A variety of *XK* mutations has been reported but no clear phenotype-genotype correlation has been found, especially for the point mutations affecting splicing sites.(16–19) Some males with the MSL also have chronic granulomatous disease (CGD). It is generally believed that patients with non-CGD McLeod may develop anti-Km but not anti-Kx, but that those with CGD McLeod can develop both anti-Km and anti-Kx.(5, 8, 20–23) but sometimes blood may not always be compatible.(24)

McLeod syndrome mimics Huntington disease; usually include progressive movement disorder (e.g: autosomal recessive chorea-acanthocytosis), cognitive impairment, and psychiatric symptoms, eventually associated with seizures, suggests that the corresponding proteins-XK.(4, 6, 9 25–30) Most importantly, given the absence of cure, it’s vital for appropriate genetic counseling, neurological examination and a cardiologic evaluation for the presence of a treatable cardiomyopathy.(5, 26, 31) Unfortunately, no clear correlations of the clinical findings with the genotype of *XK* mutations have yet been unrevealed.(32, 33)

Single-nucleotide polymorphism (SNPs) refers to single base differences in DNA among individuals. One of the interests in association studies is the association between SNPs and disease development.(34) This is the first in silico study which aim prioritize the possible mutations in *XK* gene located in coding regions and 3’UTR, to propose a modeled structure for the mutant protein that potentially affects its function. We prioritize our novel mutated SNPs for further genetic mapping studies. This will be a valuable resource for neurologists, hematologists, and clinical geneticists on this rare and debilitating disease.(35, 36)

## 2. Materials and Methods

### Data mining

The data on human *XK* gene was collected from National Center for Biological Information (NCBI) web site.(37) The SNP information (protein accession number and SNP ID) of the *XK* gene was retrieved from the NCBI dbSNP (http://www.ncbi.nlm.nih.gov/snp/) and the protein sequence was collected from Uniprot database (protein ID: 9606).(38)

### SIFT

We used SIFT to observe the effect of A.A. substitution on protein function. SIFT predicts damaging SNPs on the basis of the degree of conserved amino A.A. residues in aligned sequences to the closely related sequences, gathered through PSI-BLAST.(39) It’s available at (http://sift.jcvi.org/).

### PolyPhen

PolyPhen (version 2) stands for polymorphism phenotyping version 2. We used PolyPhen to study probable impacts of A.A. substitution on structural and functional properties of the protein by considering physical and comparative approaches.(40) It’s available at (http://genetics.bwh.harvard.edu/pp2).

### Provean

Provean is an online tool that predicts whether an amino acid substitution has an impact on the biological function of a protein grounded on the alignment-based score. The score measures the change in sequence similarity of a query sequence to a protein sequence homolog between without and with an amino acid variation of the query sequence. If the PROVEAN score ≤-2.5, the protein variant is predicted to have a “deleterious” effect, while if the PROVEAN score is >-2.5, the variant is predicted to have a “neutral” effect.(41) It is available at (https://rostlab.org/services/snap2web/).

### SNAP2

Functional effects of mutations are predicted with SNAP2. SNAP2 is a trained classifier that is based on a machine learning device called “neural network”. It distinguishes between effect and neutral variants/non-synonymous SNPs by taking a variety of sequence and variantfeatures into account. The most important input signal for the prediction is theevolutionaryinformation taken from an automatically generated multiple sequence alignment. Also structural features such as predicted secondary structure and solvent accessibility are considered. If available also annotation (i.e. known functional residues, pattern, regions) of the sequence or close homologs are pulled in. In a cross-validation over 100,000 experimentally annotated variants, SNAP2 reached sustained two-state accuracy (effect/neutral) of 82% (at anAUC of 0.9). In our hands this constitutes an important and significant improvement over othermethods.(42) It is available at (https://rostlab.org/services/snap2web/).

### SNPs&GO

Single Nucleotide Polymorphism Database (SNPs) & Gene Ontology (GO) is a support vector machine (SVM) based on the method to accurately predict the disease related mutations from protein sequence. FASTA sequence of whole protein is considered to be an input option and output will be the prediction results based on the discrimination among disease related and neutral variations of protein sequence. The probability score higher than 0.5 reveals the disease related effect of mutation on the parent protein function.(43) it’s available at (https://rostlab.org/services/snap2web/).

### PHD-SNP

An online Support Vector Machine (SVM) based classifier, is optimized to predict if a given single point protein mutation can be classified as disease-related or as a neutral polymorphism. It’s available at: (http://http://snps.biofold.org/phd-snp/phdsnp.html).

### I-Mutant 3.0

Change in protein stability disturbs both protein structure and protein function. I-Mutant is a suite of support vector machine, based predictors integrated in a unique web server. It offers the opportunity to predict the protein stability changes upon single-site mutations. From the protein structure or sequence. The FASTA sequence of protein retrieved from UniProt is used as an input to predict the mutational effect on protein and stability RI value (reliability index) computed.(44) It’s available at (http://gpcr2.biocomp.unibo.it/cgi/predictors/I-Mutant3.0/I-Mutant3.0.cgi).

### MUpro

MUpro is a support vector machine-based tool for the prediction of protein stability changes upon nonsynonymous SNPs. The value of the energy change is predicted, and a confidence score between −1 and 1 for measuring the confidence of the prediction is calculated. A score <0 means the variant decreases the protein stability; conversely, a score >0 means the variant increases the protein stability.(45) It’s available at (http://mupro.proteomics.ics.uci.edu/).

### GeneMANIA

We submitted genes and selected from a list of data sets that they wish to query. GeneMANIA approach to know protein function prediction integrate multiple genomics and proteomics data sources to make inferences about the function of unknown proteins.(46) It is available at (http://www.genemania.org/)

### Structural Analysis

#### Developing 3D structure of mutant STAT3 gene

The 3D structure of human Signal transducer and activator of transcription 3 (STAT3) protein is not available in the Protein Data Bank. Hence, we used RaptorX to generate a 3D structural model for wild-type STAT3. RaptorX is a web server predicting structure property of a protein sequence without using any templates. It outperforms other servers, especially for proteins without close homologs in PDB or with very sparse sequence profile.(47) It is available at (http://raptorx.uchicago.edu/).

#### Modeling Amino Acid Substitution

UCSF Chimera is a highly extensible program for interactive visualization and analysis of molecular structures and related data, including density maps, supramolecular assemblies, sequence alignments, docking results,conformational analysis Chimera (version 1.8).(48) It’s available at (http://www.cgl.ucsf.edu/chimera/).

#### Identification of DNA polymorphisms in miRNAs and miRNA target sites by PolymiRTS Database 3.0

We submitted SNPs located within the 3’-UTRs then the PolymiRTS checked if the SNP variants could alter putative miRNA target sites focusing on mutations that alter sequence complementarity to miRNA seed regions possibly leading to McLeod syndrome.(49) It’s available at (http://compbio.uthsc.edu/miRSNP/).

## 3. Results

**Table (1):**
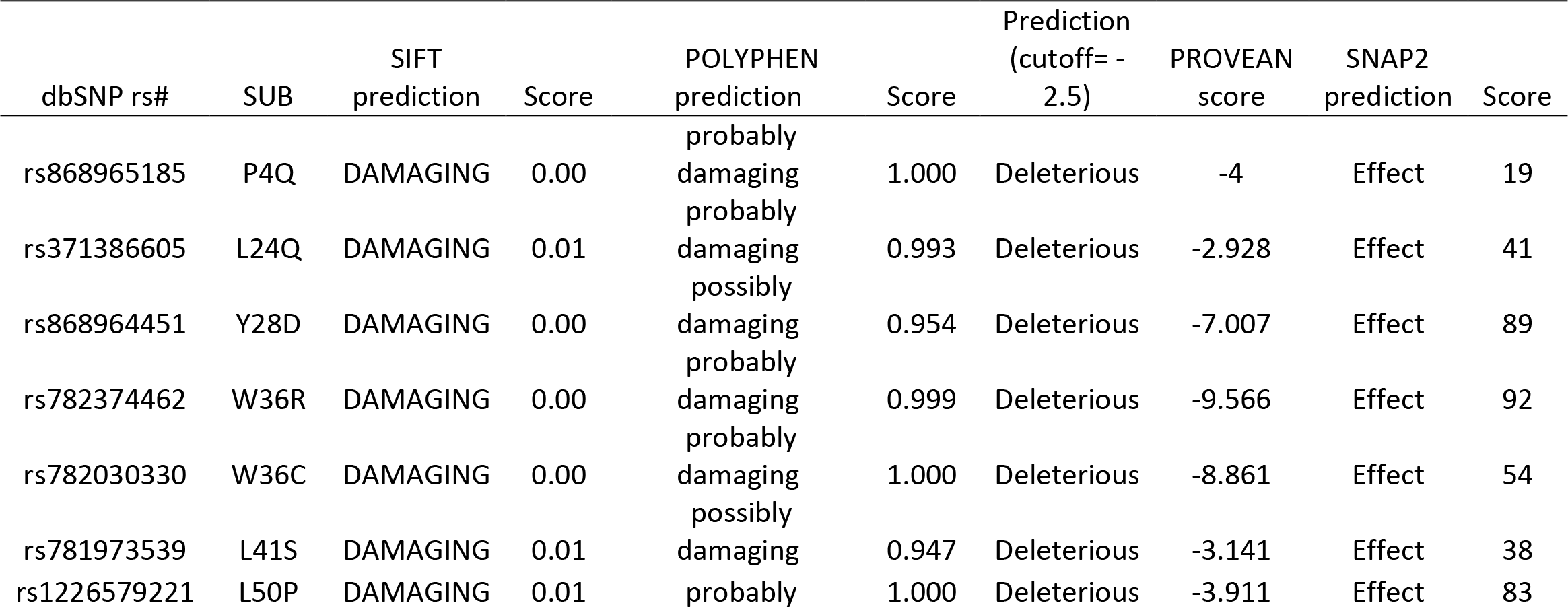
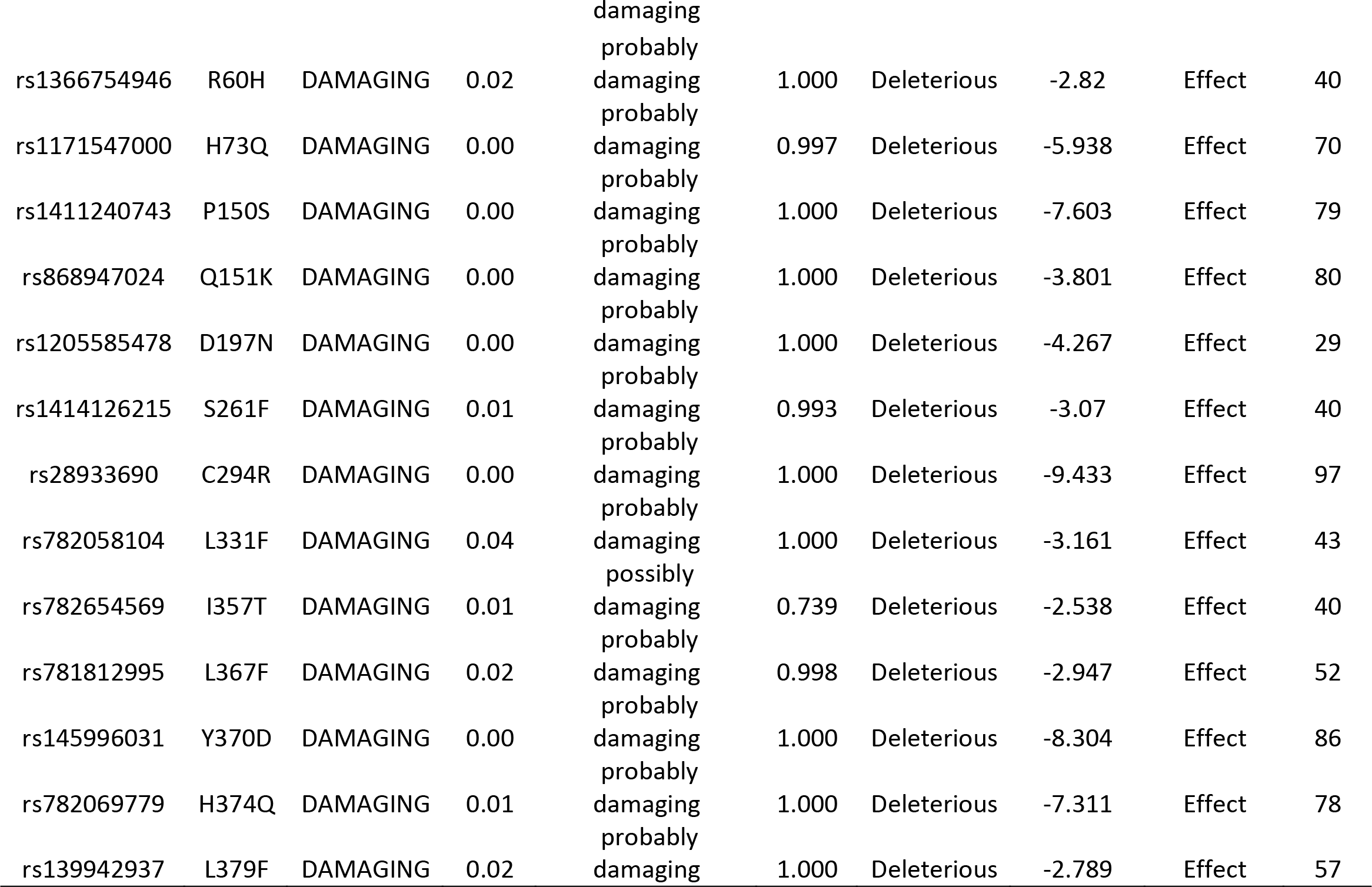
Damaging or Deleterious or effect nsSNPs associated variations predicted by various softwares:

**Table (2):** Disease effect nsSNPs associated variations predicted by SNPs&GO and PhD-SNP softwares:

| dbSNP rs# | SNPs&GO prediction | RI | Score | PhD-SNP prediction | RI | Score |
| --- | --- | --- | --- | --- | --- | --- |
| rs28933690 | Disease | 5 | 0.762 | Disease | 9 | 0.931 |
| rs145996031 | Disease | 2 | 0.596 | Disease | 7 | 0.829 |

\*RI: Reliability Index

**Table (3):** stability analysis predicted by I-Mutant version 3.0 and MUPro (also Show the 2 novel mutations):

| dbSNP rs# | Amino Acid change | SVM2 Prediction Effect | RI | DDG Value Prediction | Mupro Prediction | Score |
| --- | --- | --- | --- | --- | --- | --- |
| rs28933690 | C294R | Decrease | 2 | -0.28 | Decrease | -1.3063699 |
| rs145996031 | Y370D | Decrease | 1 | -0.9 | Decrease | -0.98561027 |

**Table (4):** The *XK* gene functions and its appearance in network and genome:

| Function | FDR | Genes in network | Genes in genome |
| --- | --- | --- | --- |
| cAMP catabolic process | 0.703371188 | 2 | 11 |
| cyclic nucleotide catabolic process | 0.703371188 | 2 | 12 |
| cyclic-nucleotide phosphodiesterase activity | 0.812112981 | 2 | 18 |
| 3',5'-cyclic-nucleotide phosphodiesterase activity | 0.812112981 | 2 | 18 |

**\*FDR:** false discovery rate is greater than or equal to the probability that this is a false positive.

**Table (5).**
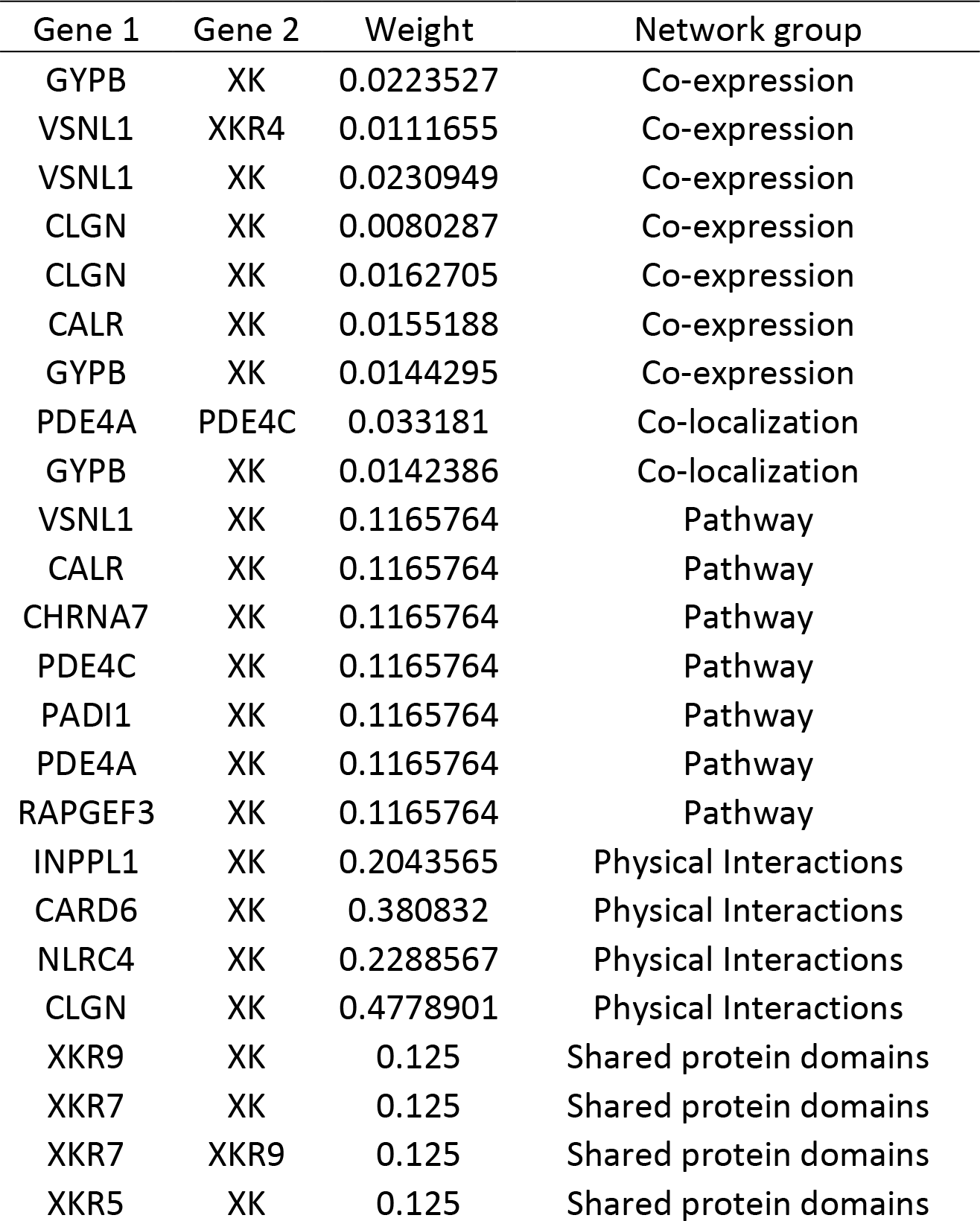
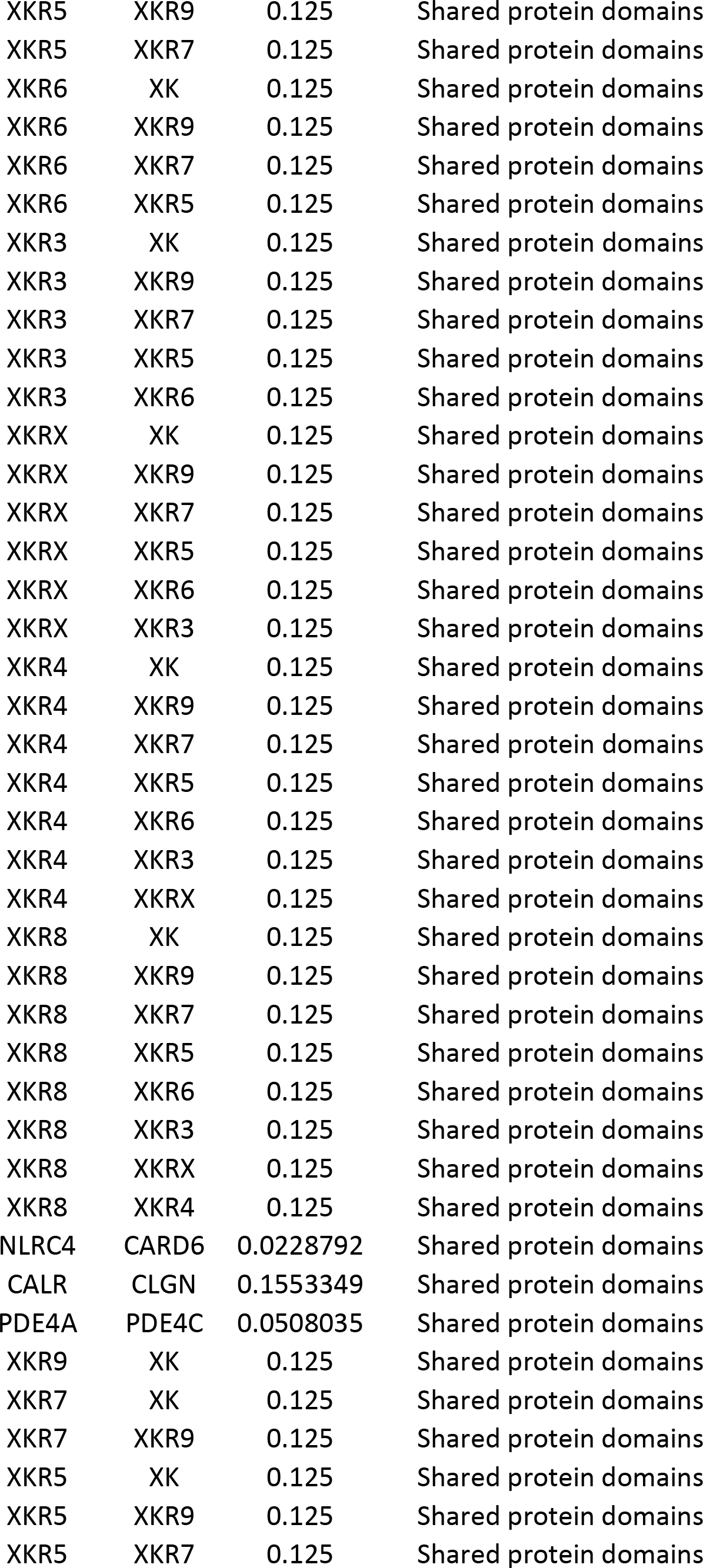
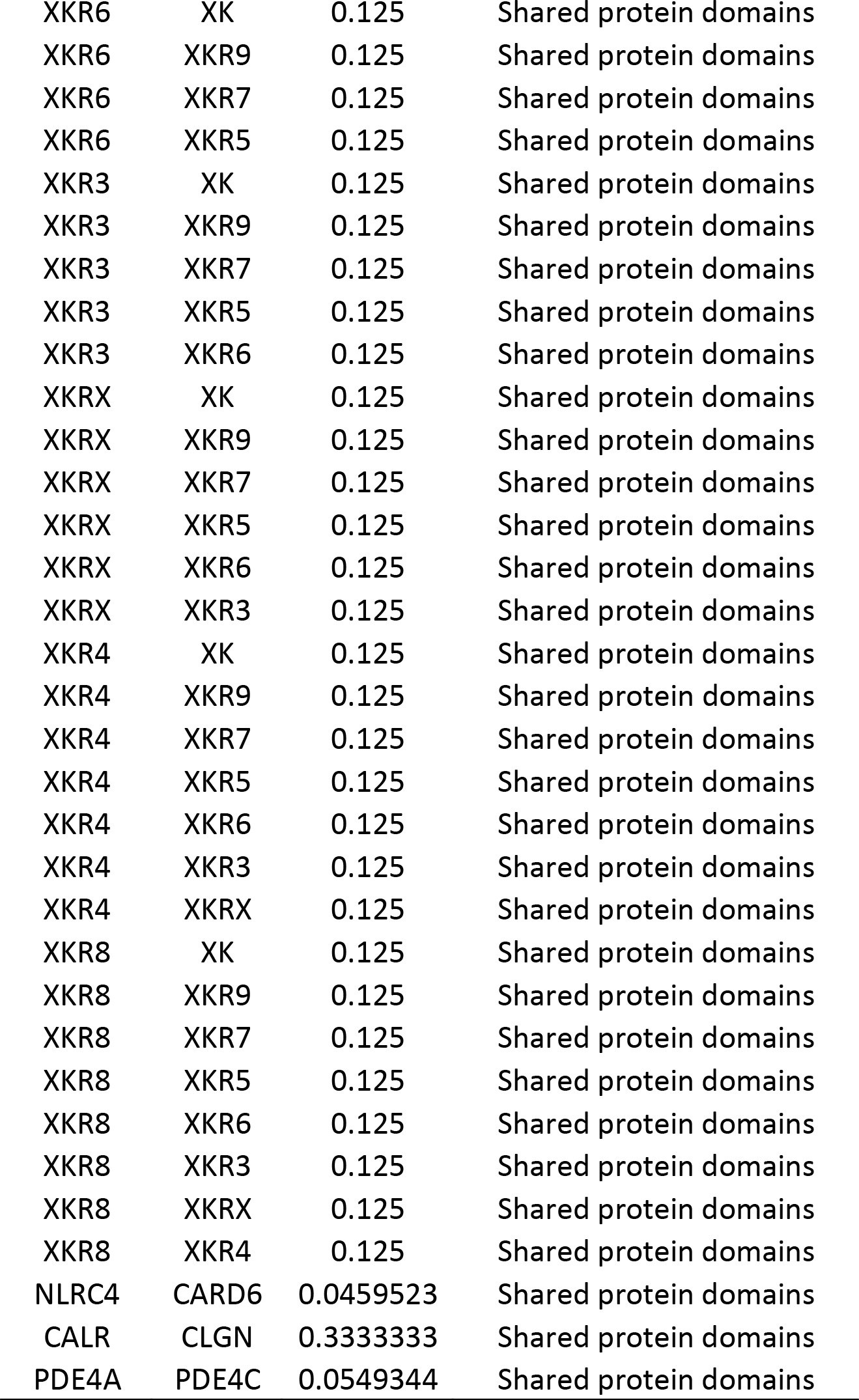
The gene co-expressed, share domain and Interaction with *XK* gene network:

**\*FDR:** false discovery rate is greater than or equal to the probability that this is a false positive. **Table (5) The gene co-expressed, share domain and Interaction with *XK* gene network:**
| Gene 1 | Gene 2 | Weight | Network group |
| --- | --- | --- | --- |
| GYPB | XK | 0.0223527 | Co-expression |
| VSNL1 | XKR4 | 0.0111655 | Co-expression |
| VSNL1 | XK | 0.0230949 | Co-expression |
| CLGN | XK | 0.0080287 | Co-expression |
| CLGN | XK | 0.0162705 | Co-expression |
| CALR | XK | 0.0155188 | Co-expression |
| GYPB | XK | 0.0144295 | Co-expression |
| PDE4A | PDE4C | 0.033181 | Co-localization |
| GYPB | XK | 0.0142386 | Co-localization |
| VSNL1 | XK | 0.1165764 | Pathway |
| CALR | XK | 0.1165764 | Pathway |
| CHRNA7 | XK | 0.1165764 | Pathway |
| PDE4C | XK | 0.1165764 | Pathway |
| PADI1 | XK | 0.1165764 | Pathway |
| PDE4A | XK | 0.1165764 | Pathway |
| RAPGEF3 | XK | 0.1165764 | Pathway |
| INPPL1 | XK | 0.2043565 | Physical Interactions |
| CARD6 | XK | 0.380832 | Physical Interactions |
| NLRC4 | XK | 0.2288567 | Physical Interactions |
| CLGN | XK | 0.4778901 | Physical Interactions |
| XKR9 | XK | 0.125 | Shared protein domains |
| XKR7 | XK | 0.125 | Shared protein domains |
| XKR7 | XKR9 | 0.125 | Shared protein domains |
| XKR5 | XK | 0.125 | Shared protein domains |
| XKR5 | XKR9 | 0.125 | Shared protein domains |
| XKR5 | XKR7 | 0.125 | Shared protein domains |
| XKR6 | XK | 0.125 | Shared protein domains |
| XKR6 | XKR9 | 0.125 | Shared protein domains |
| XKR6 | XKR7 | 0.125 | Shared protein domains |
| XKR6 | XKR5 | 0.125 | Shared protein domains |
| XKR3 | XK | 0.125 | Shared protein domains |
| XKR3 | XKR9 | 0.125 | Shared protein domains |
| XKR3 | XKR7 | 0.125 | Shared protein domains |
| XKR3 | XKR5 | 0.125 | Shared protein domains |
| XKR3 | XKR6 | 0.125 | Shared protein domains |
| XKRX | XK | 0.125 | Shared protein domains |
| XKRX | XKR9 | 0.125 | Shared protein domains |
| XKRX | XKR7 | 0.125 | Shared protein domains |
| XKRX | XKR5 | 0.125 | Shared protein domains |
| XKRX | XKR6 | 0.125 | Shared protein domains |
| XKRX | XKR3 | 0.125 | Shared protein domains |
| XKR4 | XK | 0.125 | Shared protein domains |
| XKR4 | XKR9 | 0.125 | Shared protein domains |
| XKR4 | XKR7 | 0.125 | Shared protein domains |
| XKR4 | XKR5 | 0.125 | Shared protein domains |
| XKR4 | XKR6 | 0.125 | Shared protein domains |
| XKR4 | XKR3 | 0.125 | Shared protein domains |
| XKR4 | XKRX | 0.125 | Shared protein domains |
| XKR8 | XK | 0.125 | Shared protein domains |
| XKR8 | XKR9 | 0.125 | Shared protein domains |
| XKR8 | XKR7 | 0.125 | Shared protein domains |
| XKR8 | XKR5 | 0.125 | Shared protein domains |
| XKR8 | XKR6 | 0.125 | Shared protein domains |
| XKR8 | XKR3 | 0.125 | Shared protein domains |
| XKR8 | XKRX | 0.125 | Shared protein domains |
| XKR8 | XKR4 | 0.125 | Shared protein domains |
| NLRC4 | CARD6 | 0.0228792 | Shared protein domains |
| CALR | CLGN | 0.1553349 | Shared protein domains |
| PDE4A | PDE4C | 0.0508035 | Shared protein domains |
| XKR9 | XK | 0.125 | Shared protein domains |
| XKR7 | XK | 0.125 | Shared protein domains |
| XKR7 | XKR9 | 0.125 | Shared protein domains |
| XKR5 | XK | 0.125 | Shared protein domains |
| XKR5 | XKR9 | 0.125 | Shared protein domains |
| XKR5 | XKR7 | 0.125 | Shared protein domains |
| XKR6 | XK | 0.125 | Shared protein domains |
| XKR6 | XKR9 | 0.125 | Shared protein domains |
| XKR6 | XKR7 | 0.125 | Shared protein domains |
| XKR6 | XKR5 | 0.125 | Shared protein domains |
| XKR3 | XK | 0.125 | Shared protein domains |
| XKR3 | XKR9 | 0.125 | Shared protein domains |
| XKR3 | XKR7 | 0.125 | Shared protein domains |
| XKR3 | XKR5 | 0.125 | Shared protein domains |
| XKR3 | XKR6 | 0.125 | Shared protein domains |
| XKRX | XK | 0.125 | Shared protein domains |
| XKRX | XKR9 | 0.125 | Shared protein domains |
| XKRX | XKR7 | 0.125 | Shared protein domains |
| XKRX | XKR5 | 0.125 | Shared protein domains |
| XKRX | XKR6 | 0.125 | Shared protein domains |
| XKRX | XKR3 | 0.125 | Shared protein domains |
| XKR4 | XK | 0.125 | Shared protein domains |
| XKR4 | XKR9 | 0.125 | Shared protein domains |
| XKR4 | XKR7 | 0.125 | Shared protein domains |
| XKR4 | XKR5 | 0.125 | Shared protein domains |
| XKR4 | XKR6 | 0.125 | Shared protein domains |
| XKR4 | XKR3 | 0.125 | Shared protein domains |
| XKR4 | XKRX | 0.125 | Shared protein domains |
| XKR8 | XK | 0.125 | Shared protein domains |
| XKR8 | XKR9 | 0.125 | Shared protein domains |
| XKR8 | XKR7 | 0.125 | Shared protein domains |
| XKR8 | XKR5 | 0.125 | Shared protein domains |
| XKR8 | XKR6 | 0.125 | Shared protein domains |
| XKR8 | XKR3 | 0.125 | Shared protein domains |
| XKR8 | XKRX | 0.125 | Shared protein domains |
| XKR8 | XKR4 | 0.125 | Shared protein domains |
| NLRC4 | CARD6 | 0.0459523 | Shared protein domains |
| CALR | CLGN | 0.3333333 | Shared protein domains |
| PDE4A | PDE4C | 0.0549344 | Shared protein domains |

**Table (6).**
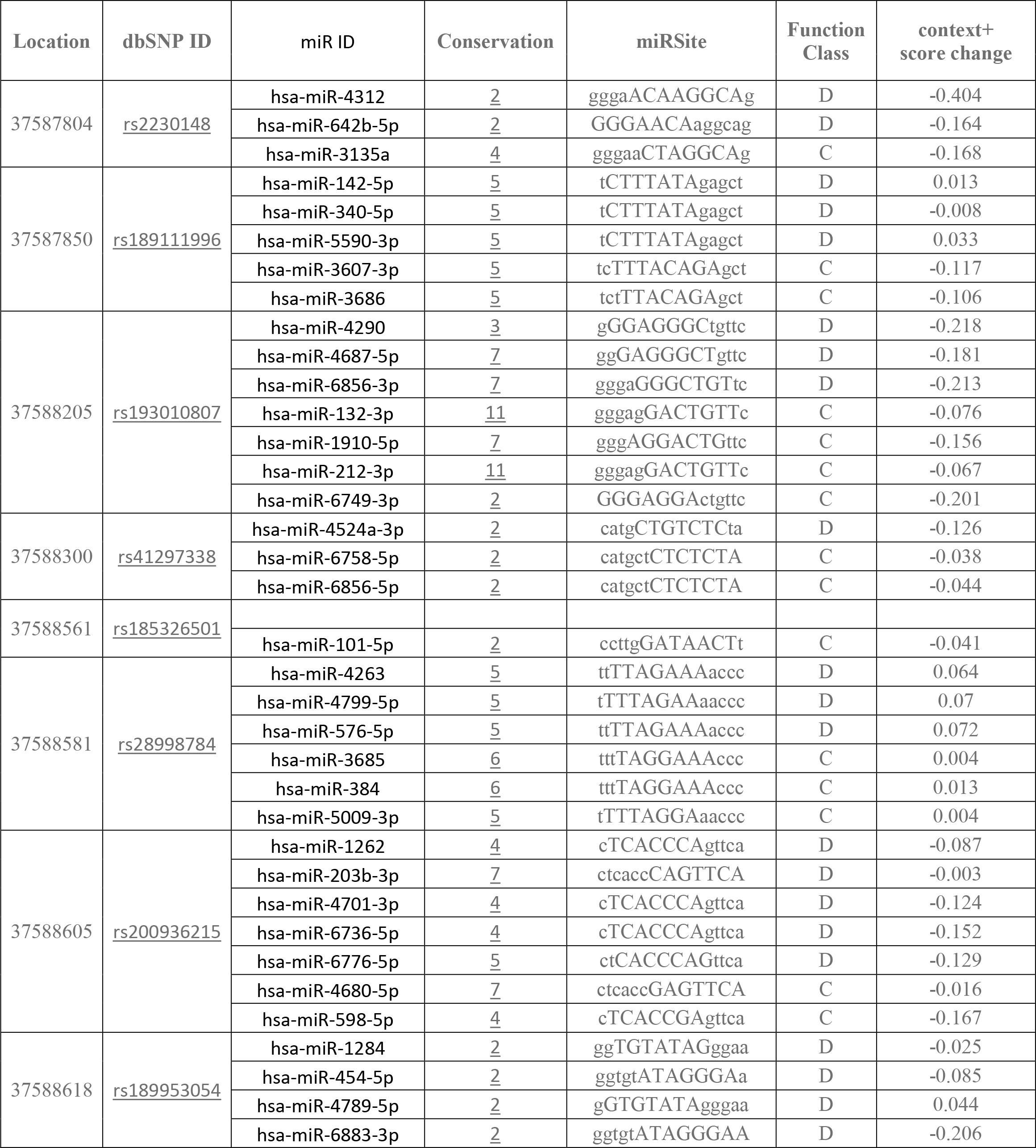
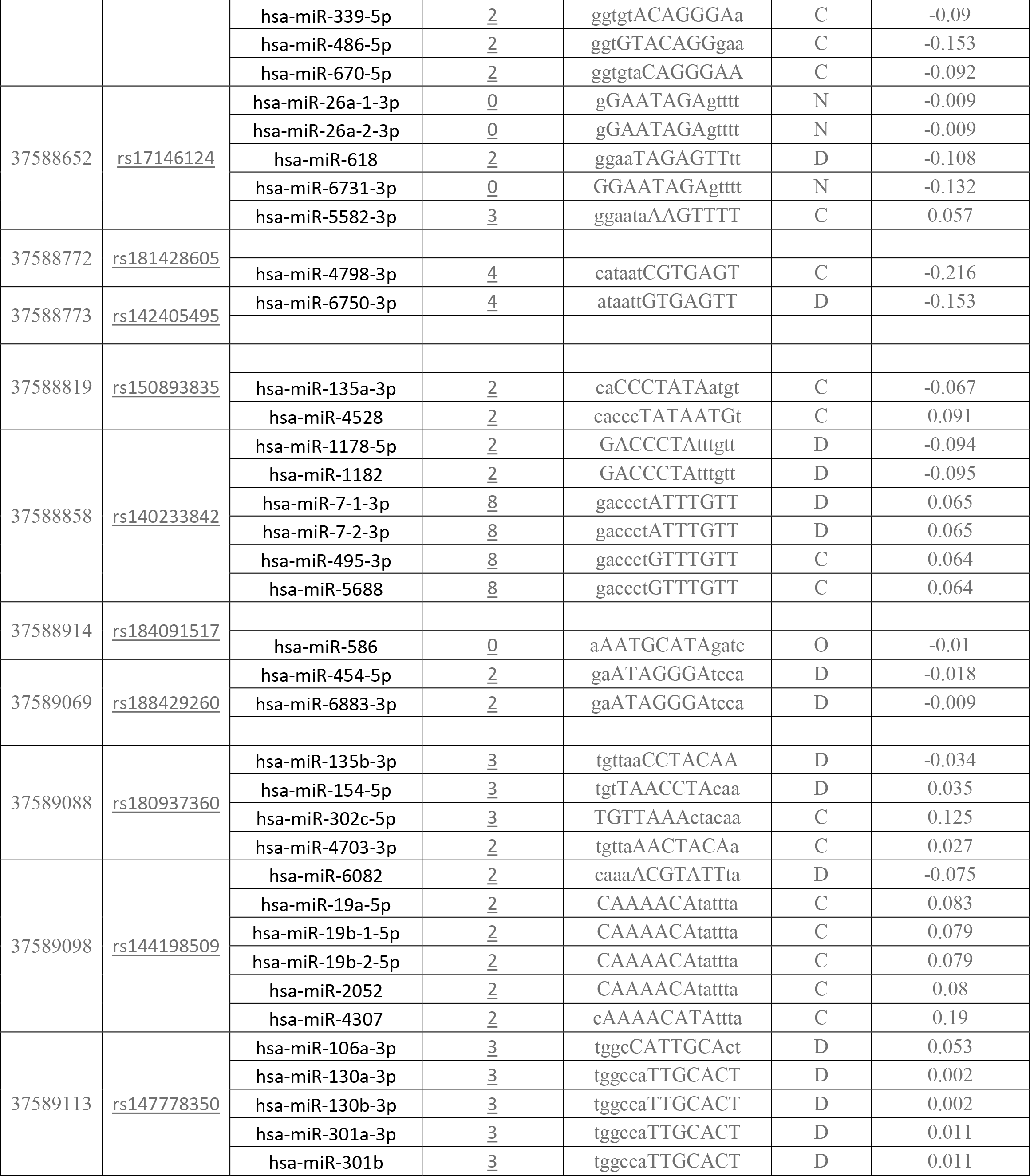
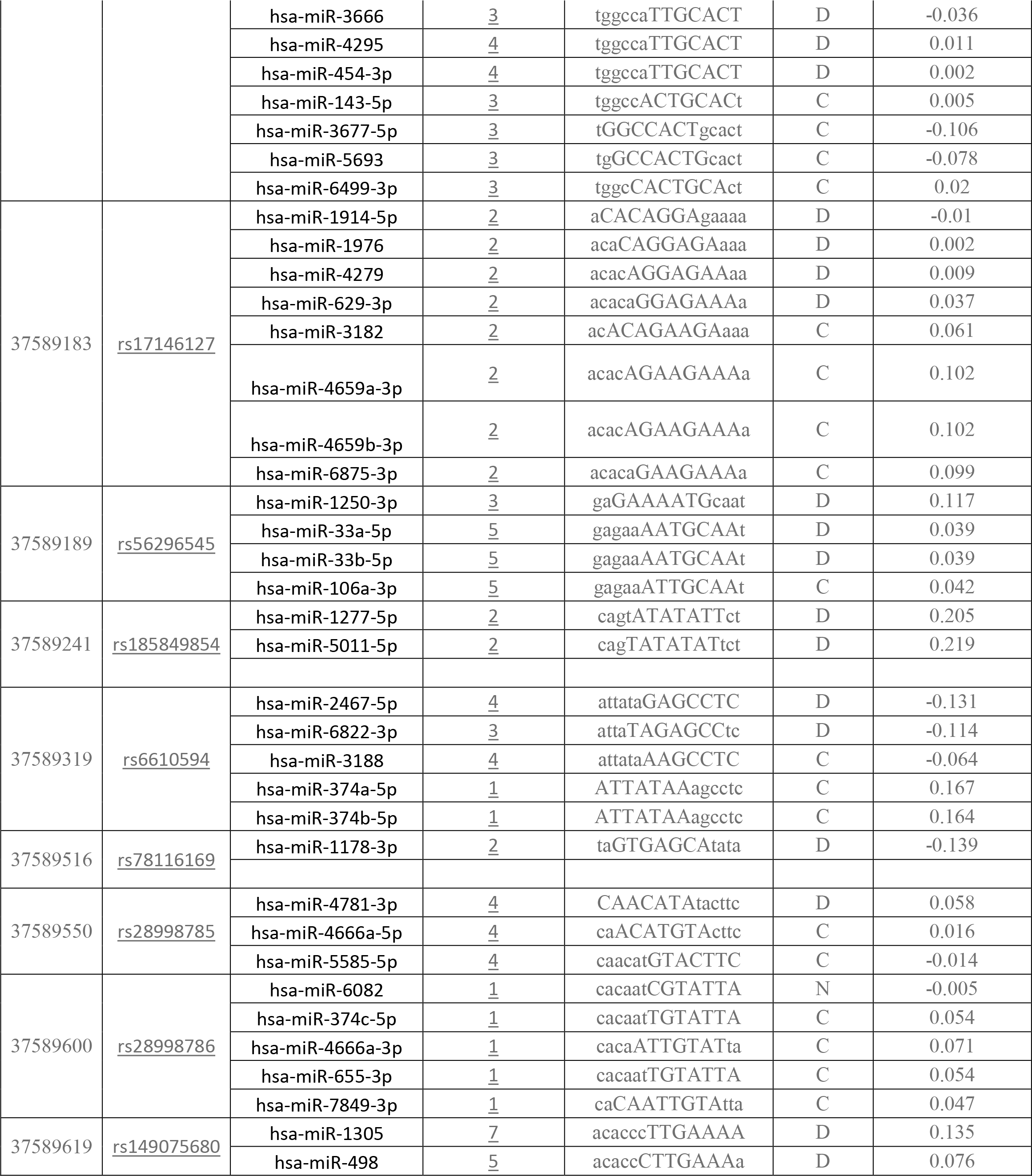
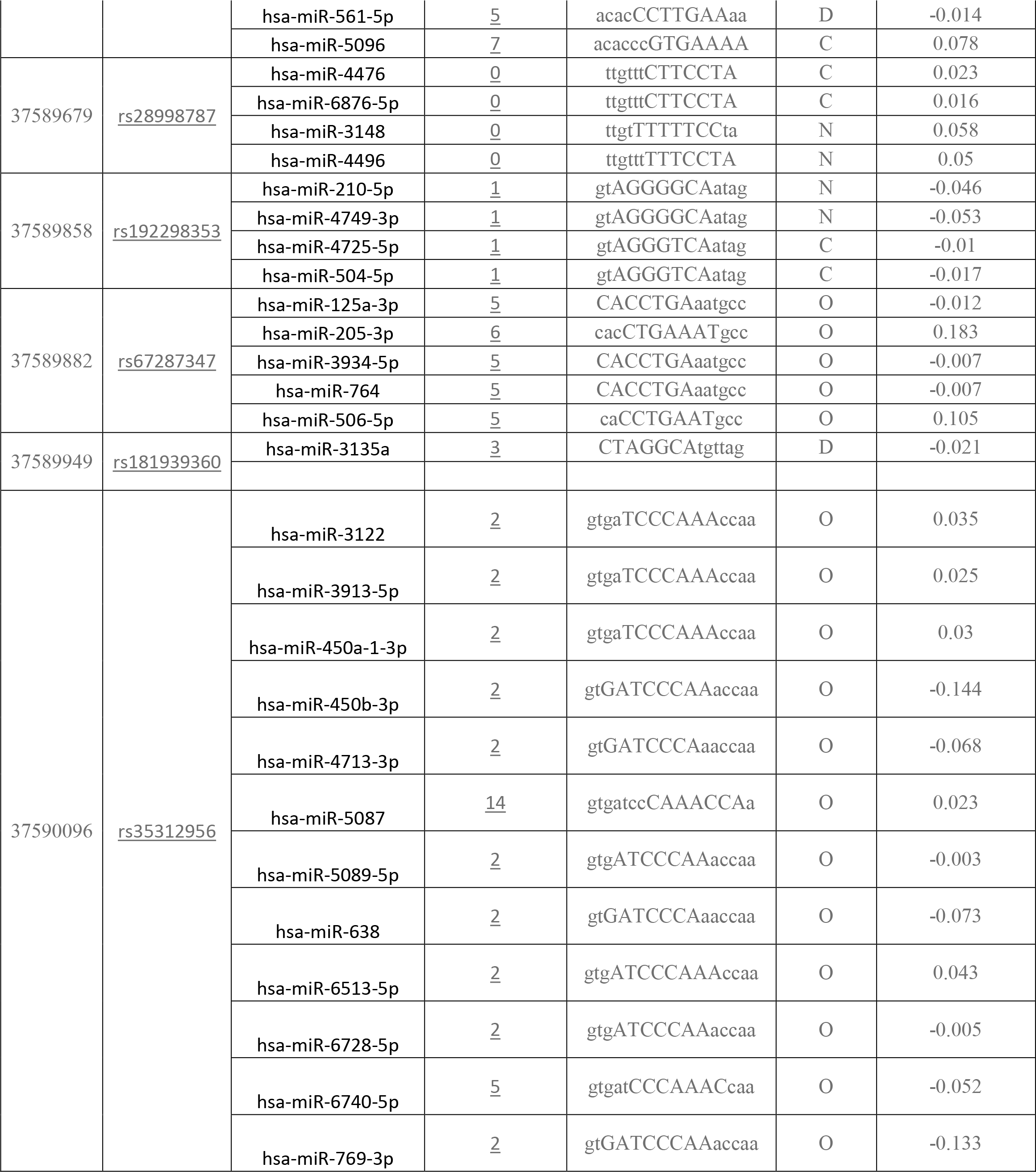
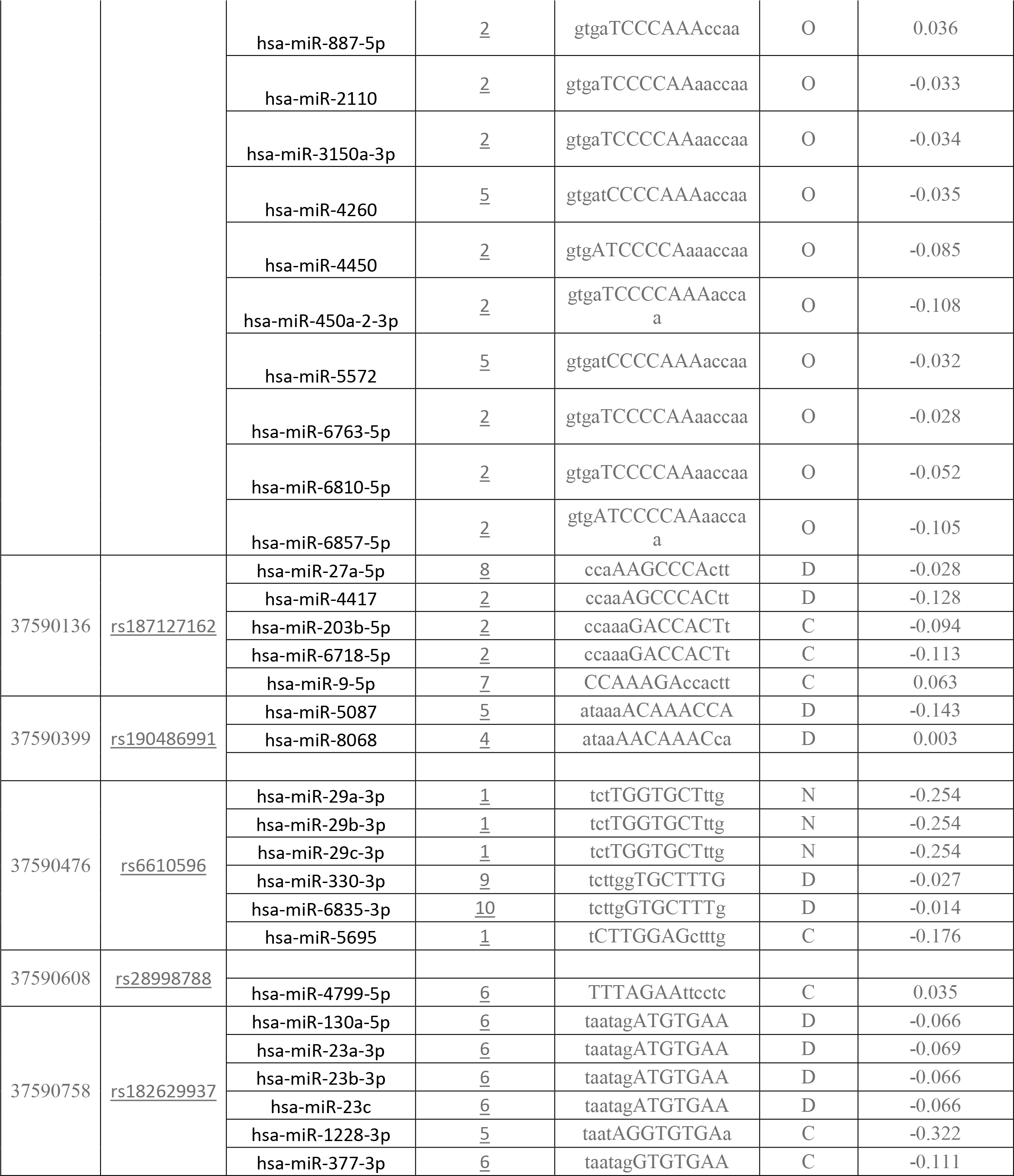
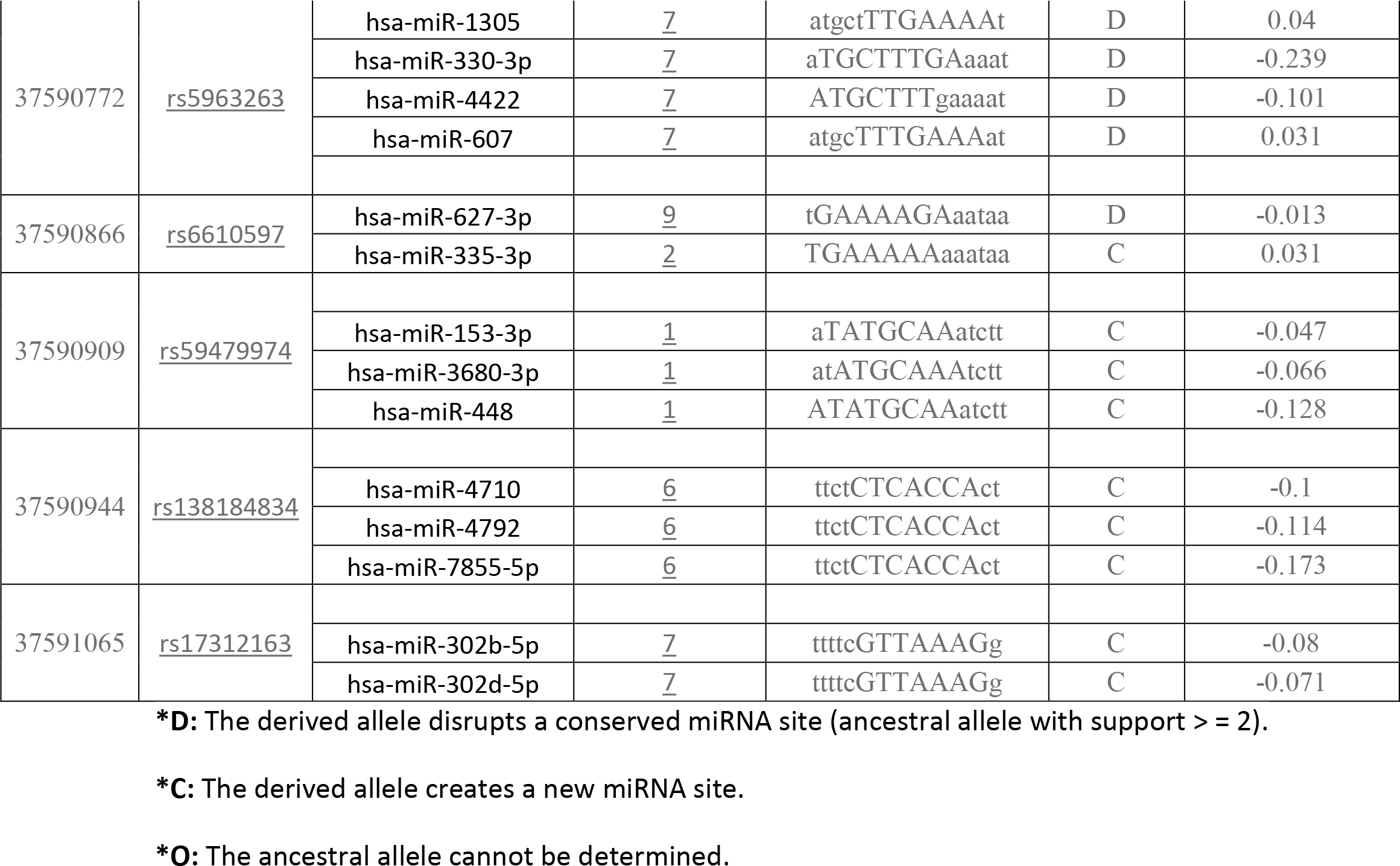
SNPs and INDELs in miRNA target sites in *XK* gene:

## Discussion

2 novel mutations have been found *that* effect on the stability and function of the *XK* gene using bioinformatics tools. The methods used were based on different aspects and parameters describing the pathogenicity and provide clues on the molecular level about the effect of mutations. Single-nucleotide polymorphism (SNP) association studies have become crucial in revealing the genetic correlations of genomic variants with complex diseases. In silico analysis has been done for many disorders for cancer related genes and other disorders (e.g: Huntington disease).(50–52)

It was not easy to predict the pathogenic effect of SNPs using single method. Therefore, multiple methods were used to compare and rely on the results predicted. In this study we used different in silico prediction algorithms: SIFT, PolyPhen-2, Provean, SNAP2, SNP&GO, PHD-SNP, P-mut, I-Mutant 3.0 and MUPro (figure 1). This study identified the total number of nsSNP in Homo sapiens located in coding region of *XK* gene, were investigate in dbSNP/NCBI Database. Out of 363 there are 104 nsSNPs, were submitted to SIFT server, PolyPhen-2 server, Provean sever and SNAP2 respectively, 35 SNPs were predicted to be deleterious in SIFT server. In PolyPhen-2 server, the result showed that 63 were found to be damaging (16 possibly damaging and 48 probably damaging showed deleterious). In Provean server our result showed that 42 SNPs were predicted to be deleterious. While in SNAP2 server the result showed that 52 SNPs were predicted to be Effect. The differences in prediction capabilities refer to the fact that every prediction algorithm uses different sets of sequences and alignments. In table (2) we were submitted four positive results from SIFT, PolyPhen-2, Provean and SNAP2 (table 1) to observe the disease causing one by SNP&GO and PHD-SNP servers. In SNP&GO and PHD-SNP servers, were used to predict the association of SNPs with disease. According to SNPs&GO and PHD-SNP revealed that, 2 and 13 SNPs were predicted to be disease related SNPs respectively. We selected the double disease related SNPs only in 2 softwares for further analysis by I-Mutant 3.0, Table (3). While I-Mutant and MUPro results revealed that the protein stability decreased which destabilize the amino acid interaction.(table3)

**Figure 1:**
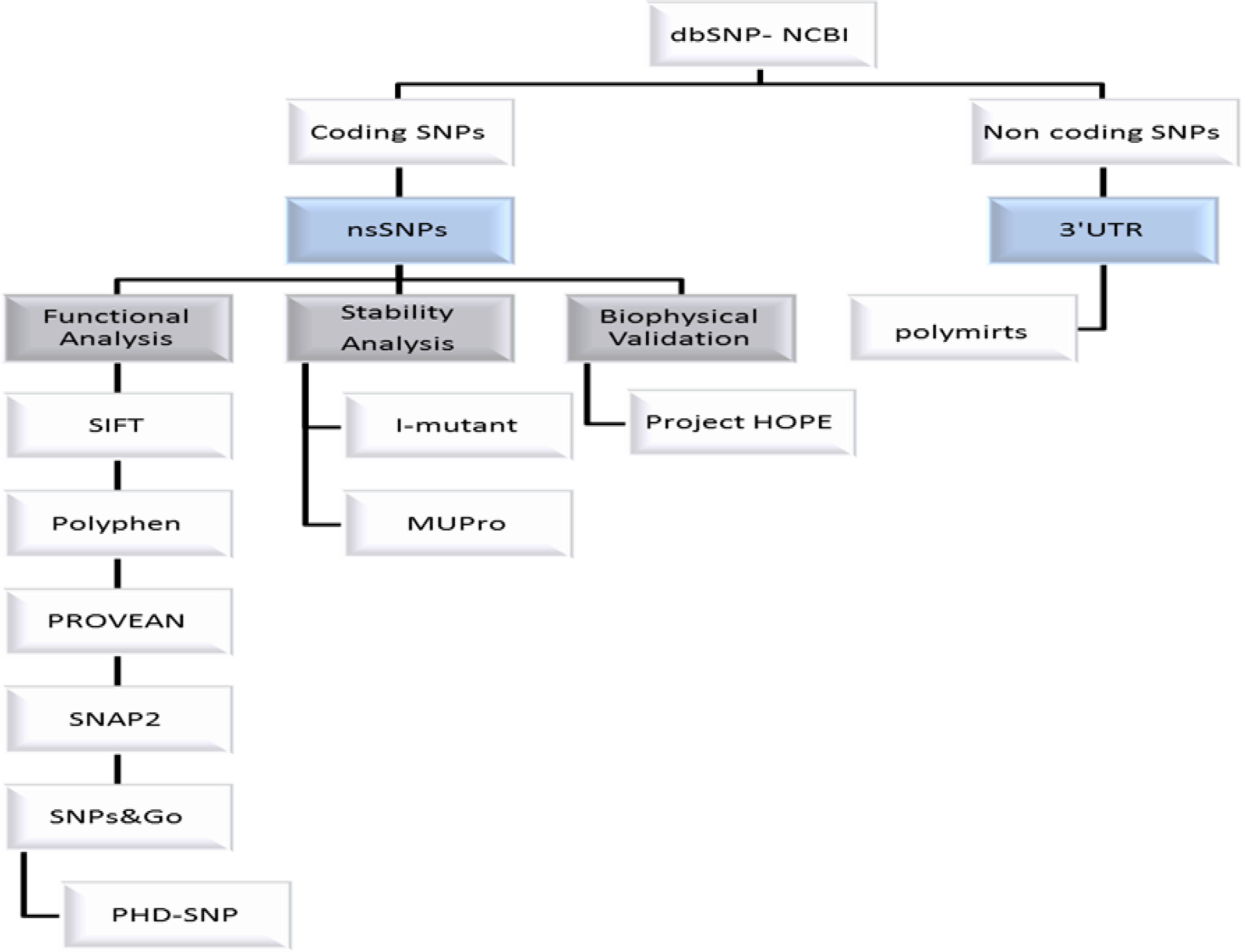
Diagrammatic representation of *XK* gene in silico work flow.

SNPs in 3’UTR of *XK* gene were submitted as batch to PolymiRTS server. The output showed that among 448 SNPs in 3’UTR region of *XK* gene, about 42 SNPs were predicted, among these 42 SNPs, 156 alleles disrupted a conserved miRNA site and 69 derived alleles created a new site of miRNA. As an example (rs200936215) SNP contained (D) allele had 5 miRSite as target binding site can be disrupts a conserved miRNA and (C) alleles had 70 miRSite disrupts a conserved miRNA site, while 27 ancestral alleles (O) cannot be determined in the all predicted 42 SNPs. Table (6) below demonstrates the SNPs predicted by Polymirt to induce disruption or formation of mirRNA binding site.

UCSF Chimera was used to visualize and analysis of molecular structures for the two deleterious nsSNPs: (rs28933690) (C294R) and (rs145996031) (Y370D). (Figures 2&3)

**Figure 2:**
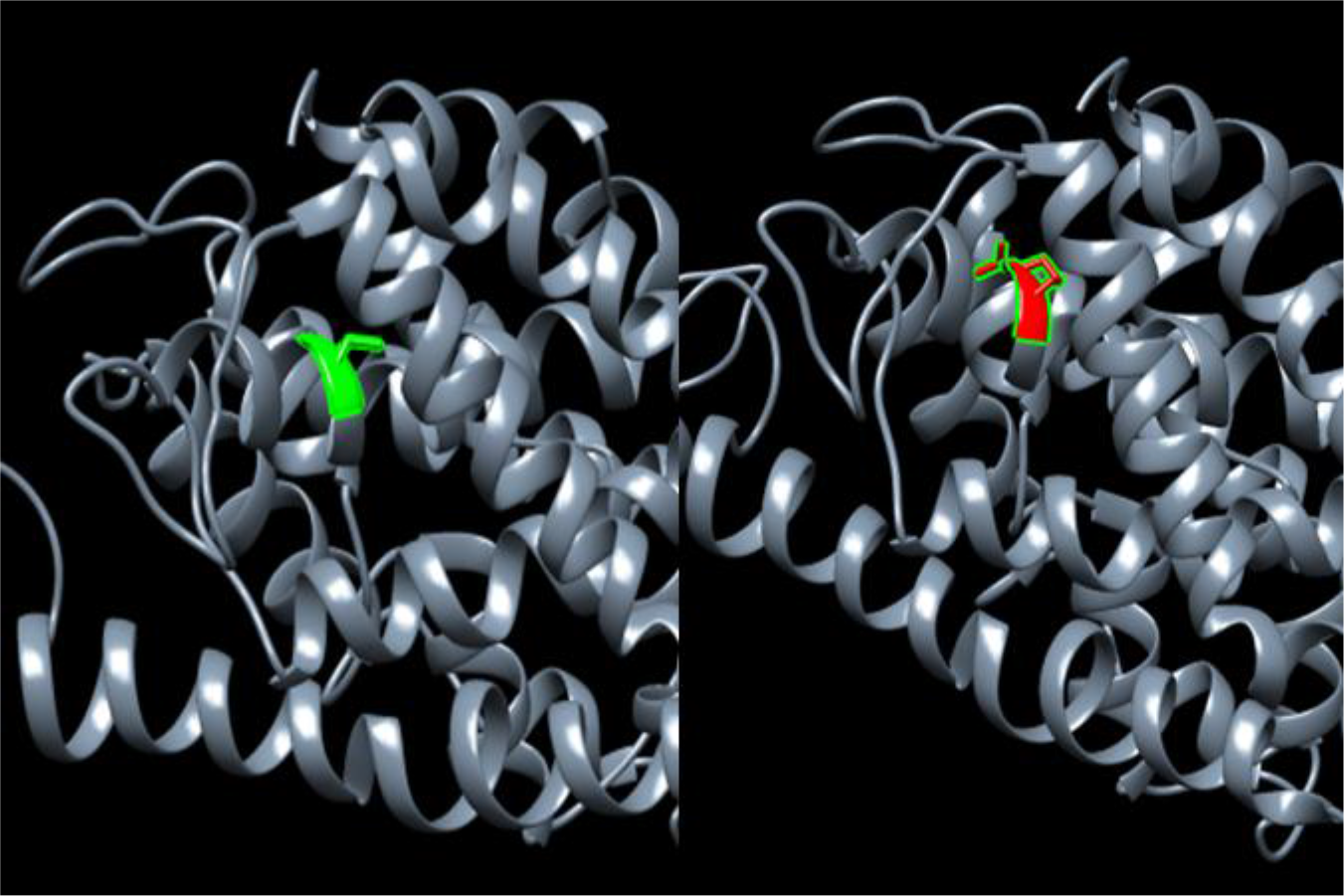
(C294R): change in the amino acid Cysteine into Arginine at position 294.

Interestingly, GeneMANIA could not predict *XK* gene function after the mutations. The genes co-expressed with, share similar protein domain, or participate to achieve similar function were illustrated by GeneMANIA and shown in figure 4, Tables (4&5).

**Figure 3:**
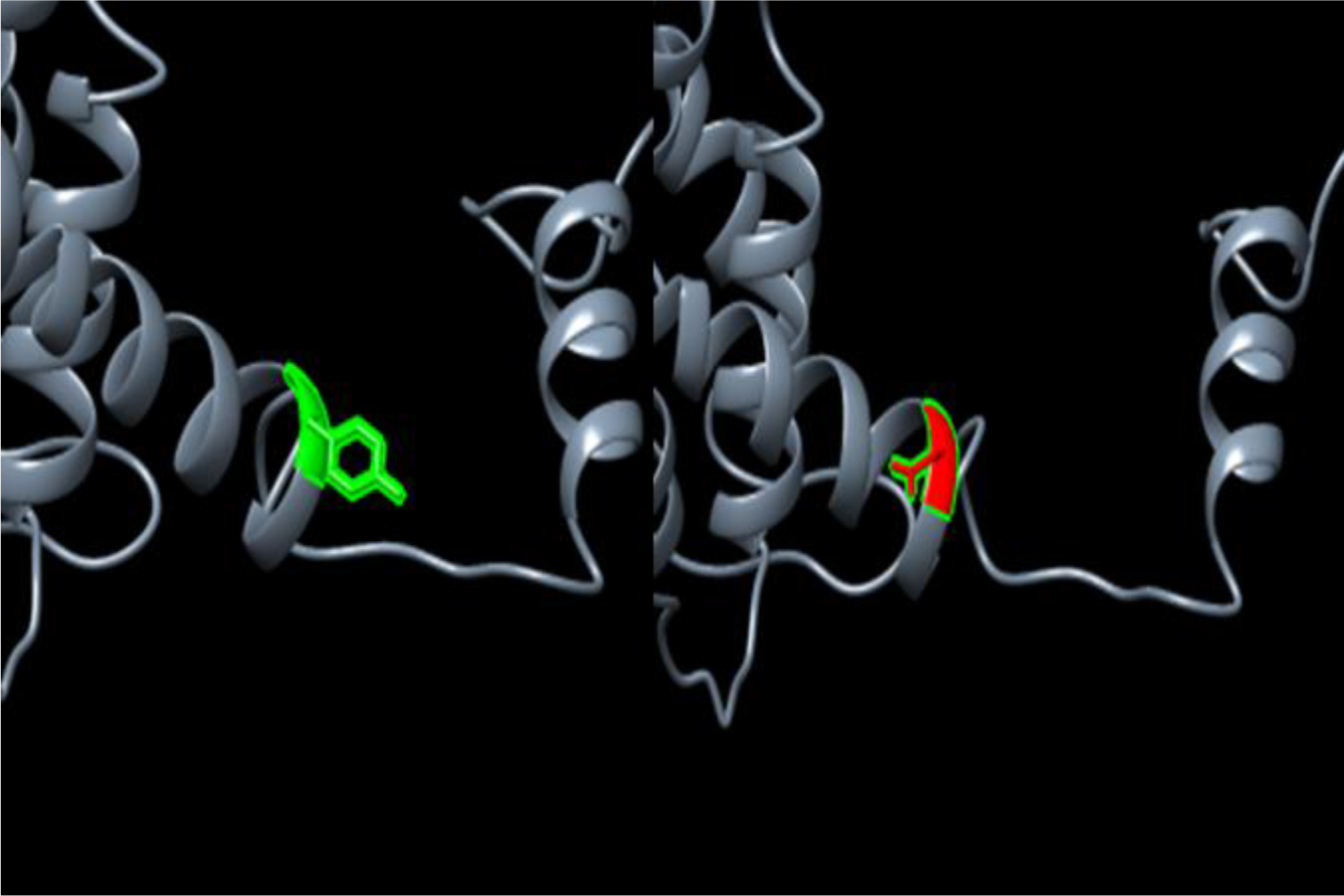
(Y370D): change in the amino acid Tyrosine into Aspartate at position 370.

**Figure 4:**
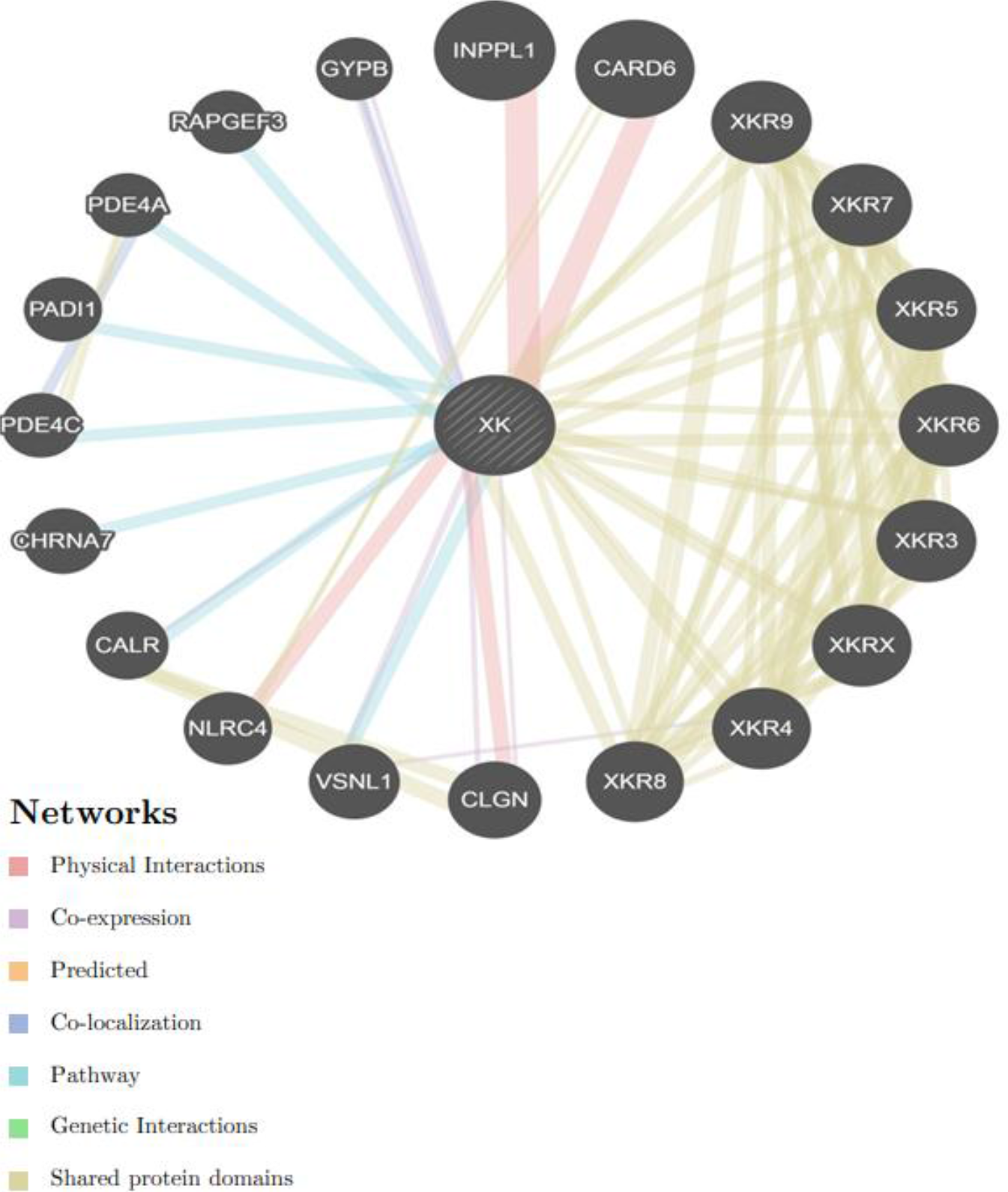
Interaction between *XK* and its related genes.

We also found that (rs28933690) (C294R) pathogenic, which is matched with the clinical result that we retrieved from dbSNPs/NCBI database. also we Retrieved (rs145996031) (Y370D) as untested we found to be damaging. Interestingly we found that, (rs145996031) (Y370D) novel SNP may also cause L1 syndrome (53).

Our study is the first in silico analysis of *XK gene* which was based on functional and structural analysis while previous studies (20, 54) based on in vivo and in vitro analysis. There is an extended phenotypic overlap between McLeod syndrome, Huntington disease and chorea-acanthocytosis, (9, 25, 29 55–57) this overlapping may help to achieve better understating for those diseases by our findings.

This study revealed 2 Novel Pathological mutations have a potential functional impact and may thus be used as diagnostic markers for McLeod neuroacanthocytosis syndrome. Constitute possible candidates for further genetic epidemiological studies with a special consideration of the large heterogeneity of *XK* SNPs among the different populations.

## Conclusion

A total of 2 novel nsSNPs were predicted to be responsible for the structural and functional modifications of XK protein These findings contribute to the understanding of the physiology of XK protein, and the pathogenetic mechanisms of acanthocytosis, myopathy, and striatal neurodegeneration in McLeod syndrome and in other associated rare neurodegenerative diseases. Also can be used for pharmacogenomics studies. Finally some appreciations of wet lab techniques are suggested to support our in silico analysis results.

## Acknowledgment

The authors wish to acknowledgment the enthusiastic cooperation of Africa City of Technology-Sudan.

